# ProFeatMap: a customizable tool for 2D feature representation of protein sets

**DOI:** 10.1101/2022.04.14.488325

**Authors:** G. Bich, E. Monsellier, G. Travé, Y. Nominé

## Abstract

**Summary:** Here, we present ProFeatMap, an intuitive Python-based website allowing to quickly display protein features such as domains, repeats, post-translational modifications location and so forth, into a highly customizable graphical 2D map. Starting from a user-defined protein list, ProFeatMap automatically extracts the main protein features from the Uniprot database. The resulting high-quality maps can help to gain insights, e.g. feature redundancy, that were previously overlooked but which may be useful for the research project. ProFeatMap is freely accessible on the web at: https://profeatmap.pythonanywhere.com/

**Availability:** Source code is freely accessible at https://github.com/profeatmap/ProFeatMap under the GPL license.

**Supplementary information:** detailed user guide of ProFeatMap

## 1. INTRODUCTION

Most -omics studies produce datasets involving substantial lists of proteins. A useful approach for examining such lists is to analyze either their protein-associated biological properties as by Gene Ontology [1], or their sequences. While Gene Ontology might be able to find over-represented features in the protein list, it loses information of their relative location, size and organization. Alternatively, sequence examination implies complex analyses as multiple alignments and requires similar proteins to be informative. An intermediate scale of analysis is to focus on features such as domains, amino-acid or domain repeats, post-translational modifications, sequence variants, secondary structures, low-complexity regions, and their organization along the sequence. Several online tools are already available to depict domain organization in a manual or semi-automatic manner (Figure 1A). However, even the most versatile programs, such as MyDomains on Prosite [2], can only scrutinize one single protein at the same time, representing a tedious and time-consuming approach even for a list containing a limited number of proteins. The interactive Tree Of Life (iTOL) tool bypasses this limitation, but specifically needs a phylogenetic tree as input [3]. To our knowledge, a versatile tool allowing feature visualization of large protein datasets in both a global and customizable way is still awaited.

**Fig. 1:**
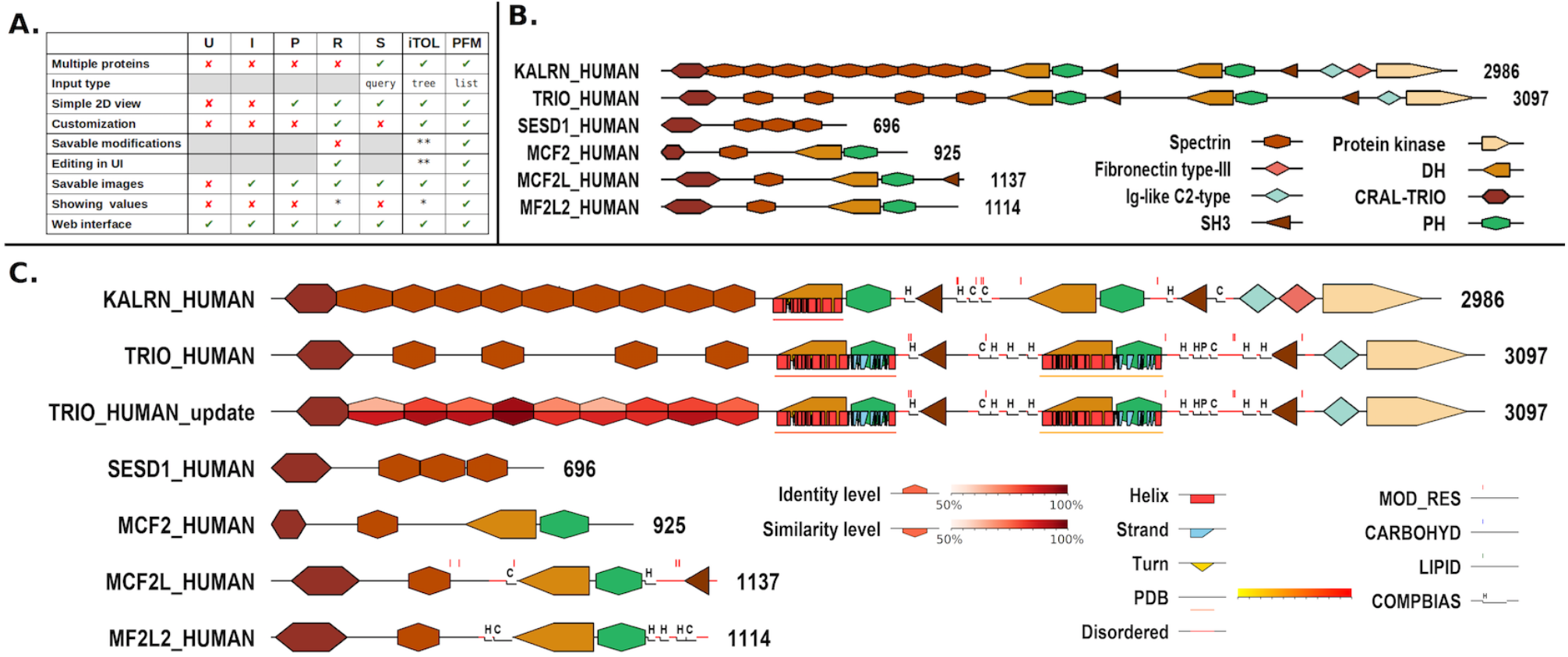
ProFeatMap overview. (A) Comparison of the main properties of different online tools that can be used for 2D protein representations. U: Uniprot; I: InterPro [5]; P:Pfam [6]; R: pRosite; S: SMART [7]; iTOL [3]; PFM: ProFeatMap. * limited possibility, ** options only accessible with an account. (B) Output of a quick ProFeatMap run. (C) Output of a ProFeatMap run when combining distinct options (same domain legends as in (B)). While Kalirin and Trio are highly similar proteins, Uniprot counts 9 spectrin repeats for Kalirin (manually curated) but only 4 for Trio (automatically curated). We performed alignments and identified in Trio the 5 missing spectrin repeats, which were subsequently added in the plot “TRIO_HUMAN_update”. The proposed spectrin repeats are colored according to sequence identity and similarity percentages obtained from the two aligned proteins. Disordered regions are highlighted in red on the sequence line. Resolved structure coverage (“PDB”) is displayed as an underlying straight line colored in heatmap mode based on the number of available structures, whereas modified residues as PTM are pointed by red marks. Composition biased regions are visualized by indents with a letter depending on the type of bias.

Here we introduce ProFeatMap, a freely available, user-friendly and interactive web interface. Gathering features directly from the Uniprot database [4], the program represents the overall domain organization and many other features in a 2D map in which the two dimensions correspond to the number and the size of the proteins contained in the list. ProFeatMap proposes default parameters for quick runs, or advanced options allowing users to produce specific plots tailored to their purposes. ProFeatMap is a powerful tool for overviewing at a glance large protein sets and gaining further insights into their common and distinctive properties.

## 2. ProFeatMap implementation

ProFeatMap is entirely coded in Python (3.9.6) [8]. It is freely accessible as a web interface developed using the Dash library (2.0.0) [9]. Therefore, it is compatible with most common web browsers. Maps and corresponding legends are drawn with the pillow library (7.2.0) [10]. The program can be run through a website without any account or registration. Alternatively, all scripts are freely accessible on GitHub and can be ran locally.

ProFeatMap follows 4 main steps in execution: protein list uploading, features extraction, addition of user numerical values (optional), and map creation and customization. A user guide for each step is found in the help sections of the web interface. A detailed user guide is available within the supplemental information.

According to a user-defined protein list, ProFeatMap automatically downloads the Uniprot file for each protein (https://uniprot.org), then extracts the protein features and subsequently determines their occurrences over the entire protein set. Are gathered: features with starting position and length, occurrence, a list of all the associated PDB files, amino acid sequences and sequence lengths. By default, ProFeatMap selects features that appear at least twice in the list.

## 3. The ProFeatMap usage

The user provides a protein list embedded into a two-columns table file (xlsx, xls, ods, csv, tsv, txt,…) containing the UniProt Accession codes and unique protein identifiers. Alternatively, a compatible list can be obtained from UniProt by downloading a protein selection through a query or the basket option. Entry lists may also be composed of various proteins obtained by experimental omics approaches, or proteins with similar segments resulting from e.g. a BLAST search. After extracting feature occurrences, ProFeatMap generates a highly customizable 2D map. An example generated for human kinases (with 489 proteins) is visible on Supplemental. This process can be done through a quick and automatic run, or by adjusting several options.

### 3.1 Quick run

In a Quick run, ProFeatMap randomly assigns a unique color and shape combination to each of the most frequent features displayed on a 2D map (Fig. 1B). The resulting map helps providing a global view of the ensemble of features. Such a quick run is preferentially used for *ab initio* lists obtained for instance in the context of omics data.

### 3.2 ProFeatMap options to highlight specific features

Alternatively, the user can select a specific feature to be highlighted in the protein map schematizing the features along the full-length protein sequences, potentially including disorder or structural information and coverage, as well as the positions of low-complexity regions or post-translational modifications. Geometrical shape and colors are also fully customizable. Optionally, the selected features can be colored according to numerical values. These could be any quantitative or qualitative values, either extracted by data mining as the number of ligands, percentage of sequence homology, reads or publications related to a given domain, or obtained experimentally as affinities, … (Fig. 1C). The same values can be used to order the proteins in the map. Another ranking can be obtained according to the feature occurrence determined in each protein, allowing to assemble proteins based on their feature content similarities. This feature is particularly useful when dealing with large lists of up to 1.000 proteins using the web interface.

### 3.3 Additional usages

ProFeatMap may help identifying recurring feature patterns in proteins. An incomplete pattern might be indicative of annotation issues in the protein sequence database. Subsequently the user may add or remove feature elements collected by further investigations directly within the web interface.

ProFeatMap also allows extracting all the sequences of a given feature in *fasta* format, or searching for motifs using regular expressions.

## CONCLUSIONS

ProFeatMap is a powerful and highly customizable tool to quickly create high quality maps displaying domains and other key features along the protein sequences for a set of proteins of interest, and potentially including quantitative or qualitative data. The general overview provided by the representation helps gaining insights into feature arrangement. The versatility and advantages of ProFeatMap can be even better appreciated when considering either a set of proteins with common features in a unique species, or a set of a particular protein originating from several species. Additionally, it is also suitable for the identification of characteristic elements such as domain repeats, that could prove to be of interest for the project.

## Supporting information

User guide for ProFeatMap

## ACKNOWLEDGMENTS

The authors thank the institutional support of the Centre National de la Recherche Scientifique (CNRS), Université de Strasbourg, Institut National de la Santé et de la Recherche Médicale (INSERM) and Région Alsace. GB is recipient of a Ph.D. grant by the Ligue contre le Cancer. The work was supported by the Ligue contre le Cancer (équipe labellisée 2015 to GT) and the Cancéropôle Grand-Est (projet Emergent 2018 to YN). The authors are grateful to Gergo Gogl for the initiation of this visualization project.

## CONFLICT-OF-INTEREST DECLARATION

The authors declare no conflict of interest.

## Notes

### Competing Interest Statement

The authors have declared no competing interest.

