## Supplementary material for "ProFeatMap: a customizable tool for 2D feature representation of protein sets": User guide for ProFeatMap

**Description**: ProFeatMap is an online web interface allowing to create highly customizable 2D representations of protein lists based on Uniprot data.

**Written by**: BICH Goran

**Contact information**:

**Last update**: March 3, 2022

**ProFeatMap version**: 1.0.0

---

**Summary**: ProFeatMap creates 2D representations (maps) of elements of interest (features) for a list of proteins based on the information available in the Uniprot database following several steps:

Step 1: The user has to input a list of proteins as Uniprot accession codes that ProFeatMap will use to download from the Uniprot database.
Step 2: Extraction of the features in the downloaded files, compiling data in a single output file.
Step 3: An optional step to add numerical values that can be shown with colormaps in Step 4.
Step 4: Creation of the map itself.

VI.2.c. Automatic feature selection 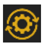 ........ 12

### I. General comments

#### I.1. Description

##### Feature

A feature is an element of interest in a protein. These features can be domains, repeats, post-translational modifications, variants, secondary structure,… These features appears in the “FT” category in Uniprot files.

##### Map

In ProFeatMap, a map is a schematic 2D representation of proteins in which features are represented according to their relative position and size on a given protein.

#### I.2. Components

##### I.2.a. Drag and Drop or Select Files

Description

ProFeatMap allow the user to upload files in several formats. It can be done either by drag and drop or by file selection.

Compatible file formats

“***.xlsx***”: 2007 and later Excel version file format.
“***.xls***”: Excel file format before 2007.
“***.ods***”: LibreOffice Calc file format.
“***.csv”***, *“****.tsv”***, *“****.tab”***, *“****.txt***”: These file format are also recognized by ProFeatMap.
Separators will be considered by descending priority order :
- tabulations
- “;” character
- “,” character

Warnings

Numerical values in columns should not be formulas and decimal separator should be “.”.

##### I.2.b. Tables

Description

Tables allow providing ProFeatMap with information needed to process the different steps. There is a total of 5 tables in the interface:

“***Protein list***”: (*mandatory*) This table contains the list of proteins that appear on the map.
“***Modifications***”: (*optional*) This table contains modifications to make during the extraction. This table is used to “correct” data or to manage more precisely the display of the proteins.

"***Numerical values***": (*optional*) This table contains the numerical values given to specific features of the proteins.

"***Shapes and colors***": (*mandatory*) This table contains the display parameters of each feature for fine tuning of the map creation. The VI.2.c. Automatic feature selection option fills up the table automatically and can be used as starting point.

"***Cut regions***": (*optional*) This table contains regions of proteins that should be hidden on the map.

<u>File name</u>

The name of the files is up to the user, only the content is used by ProFeatMap.

<u>Mandatory columns</u>

Each table has a number of columns that is recognized by ProFeatMap. Therefore, it is important to make sure all these columns names appear in the tables. Warning: Column names are case sensitive.

<u>Additional columns</u>

When building these tables, additional columns can be added. Names of additional columns must be distinct from those of mandatory columns. Spaces and special characters in the names should be avoided. Column names must be unique.

<u>Buttons</u>

Adds a row at the end of the table. Empty rows should be avoided. The new table has to be saved to be taken into account.

Used to completely clear the table, leaving only the template columns and an empty row. Clearing a table needs to be saved.

Used to save changes made in the table. Unsaved modifications will not be taken into account when executing the next step. Recovering the previous version can be done by reloading the page (unless modifications are already saved). This button will automatically be highlighted ( ) when modifications are made and switched off when clicked.

Download the latest saved table as an .xlsx file. If no table is displayed, a template file will be provided with headers only.

#### I.2.c. Map parameters buttons

<u>Description</u>

This section only applies to the three buttons under the map parameters from which the behavior is a different from the buttons below the tables.

<u>Buttons</u>

Resets all map parameters with the default values.

Saves the current selected map parameters. If ProFeatMap page is refreshed, the latest saved state will reappear. Saving is not needed to be taken into account in Step 4.

Downloads saved map parameters as an .xlsx file.

##### I.2.d. Downloadable files

Description

ProFeatMap allows the user to locally download different types of files depending on each step.

Formats

"*.xlsx*": Table, extracted data and map parameters can be downloaded as Excel files.

"*.fa*": In Step 2, sequences of a feature can be extracted. The result can be downloaded as a .fa file compatible with multiple alignment tools.

"*.png*": The maps and legend generated by ProFeatMap can be downloaded by right clicking and saving images in .png format.

### II. Quick run guide

#### II.1. Description

The quick run guide shows how to create a map by following 5 simple steps. It guides new users through the main process of creating maps while being able to customize them. It is opposed to the Map only option which directly creates a map using default options with very limited control.

#### II.2. Steps

**1.** Upload a protein list in Step 1 section in the Drag and Drop zone. (Two examples files of protein list can be downloaded for testing, TRIO and PDZ domain families)

**2.** Click on the Normal run button and wait until finished.

**3.** Go to Step 2 section and click the Run button.

**4.** Skip Step 3 and go directly to Step 4. Under the Shapes and colors table, click on the VI.2.c. Automatic feature selection and save the changes.

**5.** Click the Run button in Step 4. The map will be shown once generated.

#### II.3. Map only

Allows getting the protein map with all the default parameters. "VI.2.f. General feature parameters", "VI.2.g. Feature parameters" and "VI.2.h. Order of feature drawing" will affect the resulting map.

### III. Step 1: Protein data gathering

#### III.1. Description

The user provides ProFeatMap with a list of proteins to appear in the map. This list contains Uniprot Accession codes and names for the proteins. Subsequently, ProFeatMap downloads the Uniprot files corresponding to the provided codes.

#### III.2. Components

##### III.2.a. Protein list

Table construction

"***code***" or "***Entry***": Column containing Uniprot accession codes corresponding to the protein. The same code can be used multiple times.

"***protein***" or "***Entry name***" (optional): Column containing the names of the proteins, that will be used during all the process. It has to be unique and will serve as identifier to ProFeatMap. ProFeatMap will automatically search for Uniprot names after downloading files if names are missing and fill the table accordingly.

Uniprot search compatibility

Selected proteins of interest or basket can be exported directly from the Uniprot webpage as a compatible Excel file. "Excel" and "Uncompressed" must be selected, as shown below.

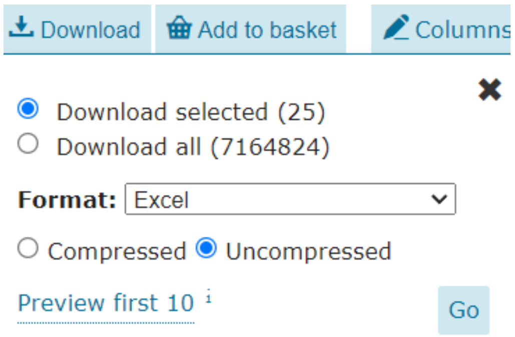

##### III.2.b. Remove organism

Description

This option removes the organism tag from the protein names in the list (i.e. TRIO_HUMAN will be transformed into TRIO). It only works if all protein names have the same tag. This feature is meant to be used on proteins names originating from Uniprot, i.e. for users interested in a single organism.

Downloading troubleshooting

If the protein list contains obsolete or invalid Uniprot codes, a section with a downloadable file containing these codes, will appear. More information on the problem can be obtained by searching for the incriminated codes on Uniprot. The codes should then be replaced or removed.

The whole protein list might show up in the file, which be most likely be a formatting error. Alternatively, a temporary loss of connection to the Uniprot database would have the same result. The current state of the connection can be obtained by refreshing ProFeatMap webpage and check the status next to the Uniprot website link:

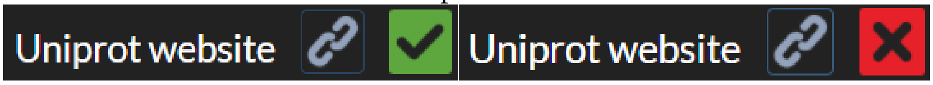

### IV. Step 2: Feature extraction

#### IV.1. Description

The main goal of this step is to extract features found in the Uniprot data files ProFeatMap gathered in Step 1. Additionally a modification table to add or remove features during the extraction process can be provided.

This step also contains additional tools: feature sequence extraction and feature/motif search by regular expression.

#### IV.2. Components

##### IV.2.a. Modification file (optional)

Description

Table filled with features to add or remove during the extraction process.

Table construction

"***ex_type***": Column filled to add ("add" or "+") or remove ("remove" or "-") a feature.

"***protein***": Name of the protein to modify.

"***feature_type***": Column to indicate the type of feature (DOMAIN, REPEAT, HELIX, BINDING, ZN_FING...). When removing a feature, the corresponding type should be checked in the output extracted data file. When adding a feature that already exists, using the same feature_type should be considered. For the addition of a not yet existing feature, DOMAIN can be used. Using other feature_type may be used for specific uses such as drawing order of features in the final map.

"***feature***": Column to indicate the name of the feature. This name will appear on the legend.

"***start***": (only for adding features) Starting position of the feature to add.

"***length***": (only for adding features) Length of the feature to add.

##### IV.2.b. Structural coverage extraction

Description

ProFeatMap searches for resolved 3D structures and calculates the number of structures available in each protein. This "coverage" can be displayed in Step 4.

Parameters

"***Step number***": (*default*: 5) The colormap is cut into a discrete series of colors. The number of resulting colors is given by the Step number. See example below.

"***Colormap thresholds***": (*default*: [1, 2, 3, 5, 10]) Each color from the discrete colormap is associated to a threshold value corresponding to the number of resolved 3D structures needed to be represented with the associated color. The color of the first step is the color at 0.2 on the base colormap. If there is only one step, the color will be the color at 1 on the base colormap. Number of structures below the first number in the array (1 in the default), will not appear on the final map.

<u>Examples</u>

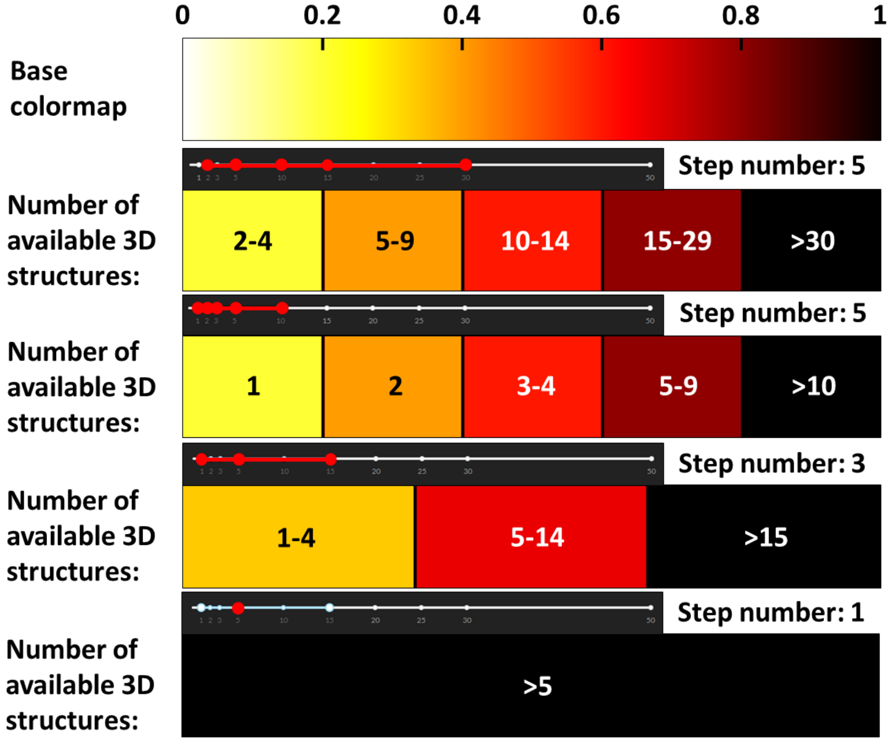

### IV.2.c. Feature sequence extraction

<u>Description</u>

Optional tool used to get the sequences of all occurrences of a given feature in the whole protein list. This tool will only appear once Step 2 has been run at least once.

<u>Parameters</u>

"***Feature name***": The name of the feature, that must correspond to the name found in the extracted data file. The name is case sensitive. With the "N-ter" and "C-ter" parameter set to 0, ProFeatMap will extract the sequences as defined in Uniprot.

"***N-ter ext***": (*default*: 0) Positive values will extend the sequence by the value towards the N-terminus of the protein. Negative values will shorten the sequence towards the C-terminus.

"***C-ter ext***": (*default*: 0) Positive values will extend the sequence by the value towards the C-terminus of the protein. Negative values will shorten the sequence towards the N-terminus.

### IV.2.d. Feature/Motif search by regular expression

<u>Description</u>

Optional tool which can be used to find the localization of features or motifs based on a regular expression. The output file can be used as modification file to add the found features during the extraction step. This tool will only appear once Step 2 has been run at least once.

Parameters

"***Feature regular expression***": To make the search needed to build a regular expression of the feature or motif. More info on how to build a compatible regular expression here: https://docs.python.org/3/library/re.html or build it directly here: https://pythex.org/

"***Feature name***": The name to give to the extracted feature/motif. This is the name that will appear on the final figure if output is added to the modification table.

##### IV.2.e. Direct download

Description

Downloads the extracted data file directly after finishing Step 2 without storing the result of the extraction in the users web browser. This option is meant to be used in case of a large lists of proteins (>1000), where ProFeatMap will most likely not be able to save the extracted data on the browser because of the lack of memory space. Activating this option will however not allow to make further steps and creating a map.

Fix options

Emptying your browsers navigation data may free enough space to be able to store the extraction. Reducing the size of the protein can be an option. If these two solutions are not working, is it always possible to install ProFeatMap locally and change the storage_type='local' to storage_type='memory' in the app.py script.

### V. Step 3: Numerical values addition

#### V.1. Description

This section can be used to add numerical values to features that can then be displayed on the map using a colorscale.

#### V.2. Components

##### V.2.a. Numerical values table

Description

A table containing a list of features and the numerical values that should be associated.

Table construction

"***protein***": name of the protein that contains the feature.

"***feature***": name of the feature. This name has to correspond to the exact name of the feature that appears in the extracted data file.

"***start***": the starting position of the feature. This value has to appear in the extracted data file. It is used to identify the feature (if multiple).

"***condition_x***": multiple conditions can be added. One column per condition is needed and must have a unique user defined name. Spaces should be avoided. These conditions can then be selected in Step 4 for the map creation.

Values

The values given by the user should be normalized (between 0 and 1). -1 will be interpreted as missing value and will appear in grey on the map. No value will result in the use of the specified “color” and “contour_color”.

### VI. Step 4: Map creation

#### VI.1. Description

Creates the map of the protein list using shapes and colors either defined automatically or defined by the user potentially considering cut regions. Map parameters are used to change the overall map generation, the display of specific features, the order of feature drawing and the condition (if any) to show numerical values.

#### VI.2. Components

##### VI.2.a. Shapes and colors table

Description

The shapes and colors table contains the list of features to appear as specific shapes and colors on the protein map.

Table construction

“***shape***”: shape of the feature. A list of available shapes can be found below the table.

“***orientation***”: some shapes have associated orientation that have to be specified.

“***height***”: (*default*: vertical stretch factor) corresponds to the height, in pixels of the feature.

“***contour_color***”: (*default*: black) color of the contour of the shape.

“***contour_colormap***”: (*default*: none) used to show numerical values on the contour of the shape. If the Uniform shape fill/contour is selected, a “colormap” is defined and no “contour_colormap” is selected, the edges share the “colormap” with the inside of the shape.

“***contour_threshold***”: (*default*: none) Only used if a “contour_colormap” is defined. A value between 0 and 1 can be indicated. Values below this threshold will appear in the color defined in “contour_color”. Values above the threshold will use the “contour_colormap”.

“***color***”: (*default*: white) color used to fill the shape.

“***colormap***”: (*default*: none)

“***threshold***”: (*default*: none) Only used if a colormap is defined. A value between 0 and 1 can be indicated. Values below this threshold will appear in the color defined in “color”. Values above the threshold will use the “colormap”.

“***pensize***”: (*default*: protein thickness) corresponds to the thickness of the contour of the feature.

Showing other features

Adding “Others” as feature name will show all features that are not listed in the table using a white rectangle with black contour by default.

#### VI.2.b. Protein cuts table

Description

To shorten specific proteins by hiding a fraction of the protein (indicated by a "//" on the map) and all included features.

Table construction

"***protein***": Name of the protein to be cut out.
"***start***": Starting position of the region to hide.
"***length***": Length of the region to hide.

#### VI.2.c. Automatic feature selection

Description

Automatic feature selection will search in the extracted the features either the most represented features or features that appear more than a given occurrence in the list. Only features under the DOMAIN, REPEAT, MOTIF or REGION tag are considered. If the feature is one of the most common ones (DOMAIN and REPEAT in the human proteome), the default shape and color will be used. Other features will have a random shape and color. All features that are not represented enough will be represented by the "Others" category. By default "more than 2 occurrences" is used.

Parameters

"***x most represented features***": (*default*: square root of the number of proteins) automatic feature selection will choose a shape and color for the top x most represented features
"***more than x feature occurrences***": (*default*: 2) automatic feature selection will choose a shape and color for features that have been found at least x times in the protein list

Lock seed

This option can be toggled to fix the current seed used for random picking of shapes and colors by the automatic feature selection.

#### VI.2.d. Sorting

Description

ProFeatMap features several sorting options impacting the order of appearance of the proteins on the map.

Options

"***None***": order of the proteins as defined in the table in Step 1.
"***abc***": to sort the protein list in alphabetical order.
"***feature_number_distance***": (*default option*) to let ProFeatMap sort automatically the proteins by gathering those with similar feature content. WARNING : This sorting is quite intensive. It should not be used with large lists of proteins **(>2,000)**. The resulting sorting can be saved with the "Latest sorting protein list" button. This list can be used as input list, and None put as sorting. This way of proceeding is strongly advised for lists with more than 200 proteins in order to decrease the drawing process time.
"***value***": see VI.2.e. Value related below

#### VI.2.e. Value related

Description

Sorting by value is only available if the user has given numerical values in Step 3. This sorting will order proteins by descending values. Each protein is represented by the highest value if multiple occurrences of the target feature is found.

Parameters

"***Case to draw***": (*default*: None) all conditions defined in the numerical value file will appear in this list when Step 3 is run. Selecting a condition will affect the values displayed on the map.

"***Focus on***": (*default*: None) the name of the feature by which the map should be sorted must be inputted here.

"***Threshold***": (*default*: None) by indicating a threshold (float value), allows to remove from the map all proteins with focused on feature with values below the threshold.

#### VI.2.f. General feature parameters

Description

General feature parameters will affect general visual aspects of the map such as horizontal and vertical stretch, protein thickness, text sizes and the display of the protein length.

Parameters

"***Horizontal stretch factor***": (*default*: 1) multiplying factor of the horizontal length on the map. > 1 will cause the proteins and features to appear longer, whereas < 1 will make them appear smaller. Support floats.

"***Vertical height***": (*default*: 20) vertical space used by each protein. Increasing the value will make the figure bigger. It will also cause all features shapes that have no "height" specified to have this value as default value.

"***Protein thickness***": (*default*: 3) thickness of the line representing the protein. This value is the default pensize value for feature shapes (contours), if none is specified.

"***Protein name size***": (*default*: 30) size of the text showing the protein names.

"***Biased regions text size***": (*default*: 20) size of the text shown above the composition biased regions.

"***Show protein length***": (*default*: y) toggle to activate or deactivate the display of the protein length next to each protein.

"***Consistent shape fill/contour***": (*default*: n) toggle to activate or deactivate the display the standardization of the filling color and the contour color. When activated, all feature shapes that have no "contour_color" specified, will appear the same color as the filling color (if there is one). It also applies to colormaps.

#### VI.2.g. Feature parameters

Description

These parameters can be toggled on or off to make specific features appear or disappear from the created map.

Parameters

"***3D structure coverage***": (*default*: n) As defined during extraction (see Structural coverage extraction), toggling this option will show a line where resolved 3D structures have been found in the PDB database. The line is colored depending on the number of structures.

“***Secondary structure***”: (*default*: n) Secondary structures (helixes, strands and turn) will be represented on the proteins.

“***Disorder***”: (*default*: y) When toggled, predicted disordered regions will be shown on proteins.

“***Modified residues***”: (*default*: n) When toggled, modified residues (phosphorylation, glycosylation,…) will appear on proteins.

“***Composition biased regions***”: (*default*: None) Selected biases can be selected, and will appear on the proteins.

Feature parameters default

Clicking this button will add to the current shapes and colors table the default representation of each selected “Feature parameter” in the “Map parameters”. The addition of these representations will overwrite the default representation. This will also cause these parameters to be always displayed whenever the corresponding parameter is toggled or not.

#### VI.2.h. Order of feature drawing

Description

This list of feature categories defines the order in which the features will be drawn on the proteins. In case of overlapping features, the category appearing the latest in the list will appear above. Default parameters should avoid most overlaps.

Parameters

(*default*: DISORDER, DOMAIN, CHAIN, INIT_MET, ZN_FING, DNA_BIND, REGION, ACT_SITE, METAL, SITE, LIPID, HELIX, STRAND, TURN, CONFLICT, CARBOHYD, BINDING, MOTIF, MOD_RES, COMPBIAS, REPEAT, VARIANT, PDB)
